# Is testosterone responsible for athletic success in female athletes?

**DOI:** 10.1101/557348

**Authors:** Ildus I. Ahmetov, Thomas R. Roos, Albina A. Stepanova, Elnara M. Biktagirova, Ekaterina A. Semenova, Irina S. Shchuplova, Larisa V. Bets

## Abstract

The aim of this study is to determine the interrelationship between the resting serum testosterone (T) levels (an inherited trait) of female athletes from different types of sporting events and their athletic success. The study involves 599 Russian international-level female athletes (95 highly elite, 190 elite, and 314 sub-elite) and 298 age-matched female controls. All subjects were age 16-35 years old and to the best of our knowledge have always tested negative for performance enhancing substances. The athlete cohort was stratified into four groups according to event duration, distance, and type of activity: 1) endurance athletes, 2) athletes with mixed activity, 3) speed/strength athletes, and 4) sprinters. Athletic success was measured by determining the level of achievement of each athlete. The mean (SD) T levels of athletes and controls were 1.65 (0.87) and 1.76 (0.6) nmol/L (*P*=0.057) with ranges of 0.08-5.80 and 0.38-2.83 nmol/L in athletes and controls, respectively. No significant differences in T levels were found between different groups of athletes. T levels were positively correlated (*r*=0.62, *P*<0.0001) with athletic success in sprinters (runners, cyclists, kayakers, speed skaters, swimmers). Moreover, none of the sub-elite sprinters had T > 1.9 nmol/L, while 50% of elite and highly elite sprinters had T > 1.9 nmol/L (95% CI: 2.562-862.34; OR=47.0; *P*<0.0001). We do not observe the benefits of having high T levels for success in other groups of athletes. Conversely, highly elite middle-distance (*P*=0.235) and mixed activity athletes (*P*=0.096) tended to have lower T levels than less successful athletes. Our data suggest that the measurement of the serum T levels significantly correlates with athletic success in sprinters but not other types of athletes and in the future may be useful in the prediction of sprinting ability.

## INTRODUCTION

Testosterone (T), which is normally present in the circulation of both men and women, has an anabolic effect and stimulates the growth of muscle by increasing muscle protein synthesis and inhibition of protein breakdown via a ubiquitin-mediated pathway^1^. There is agreement that T supplementation increases muscle mass and maximal voluntary strength and reduces body fat^2–4^. High T concentrations or precursor androgens were found to be associated with greater lean body mass and higher bone mineral density in women^5–7^. T also acts on specific substrates in the brain to increase aggression and motivation for competition^8,9^. Thus, high endogenous T concentrations may confer both psychological and physiological advantage in sports^4^.

Exercise (especially resistance exercise training) is globally able to induce an increase in circulating T and other hormones^10^. Therefore, it was suggested that the measurements of T and T to cortisol ratio may be useful biological markers of training responses^11,12^. Due to the dynamic regulation of endogenous T production, including the acute effects of competition and exercise, T concentrations may vary considerably within and among individuals^4^, which can be determined, in part, by genetic polymorphisms. Studies in at least male twins indicate that there is a strong heritability of serum T, with genetic factors accounting for 65% of the variation in serum T^13^. Several polymorphisms of the sex hormone-binding globulin (*SHBG*; transports androgens and estrogens in the blood), α-actinin-3 (*ACTN3*; interacts with calcineurin which negatively regulates the production of T), aromatase (*CYP19A1*; converts androgens to estrogens) and other genes were identified to be associated with serum T concentration in different populations including athletes of both sexes^14–16^. Given an important physiological role of T in the growth and metabolism, one might suggest that natural T levels could predict athletic potential of an individual and thus the measurement of serum or salivary T may be used in sports selection. On the other hand, other factors may influence T concentrations in female athletes, including circadian variation, oral contraceptive use, nutrition, load demands of training etc. ^17^.

There is much debate over whether or not women whose testosterone (T) levels cross into the male range (i.e. ≥ 10 nmol/L) should not compete with other women unless it was shown that they are resistant to the effects of T (androgen insensitivity). Recent study has shown that female athletes (*n* = 300) with high T levels have a significant competitive advantage over those with low T in 400-800 m running, hammer throw and pole vault^18^. On the other hand, several specialists in this area argue that the IOC and the IAAF policies should be withdrawn on the grounds that they are discriminatory towards women and place some female athletes at risk of unnecessary and potentially harmful investigations and treatments^19,20^. The aim of the study was to determine the interrelation between the resting serum T levels of female athletes from different sporting disciplines and their athletic success.

## MATERIALS AND METHODS

### Subjects

The study involved 599 Russian female athletes (age: 16-35 years) from the National teams who were tested negative for doping substances. The athletes were stratified into four groups according to event duration, distance and type of activity. The first group (‘endurance group’; *n* = 133) included long (*n* = 77) and middle (*n* = 56) distance athletes (31 rowers, 6 800-10000 m runners, 18 biathletes, 5 middle-distance kayakers, 8 1.5-3 km speed skaters, 38 cross-country skiers, 1 marathon runner, 10 200-10000 m swimmers, 13 triathletes, 3 middle-distance short-trackers). The second group (‘mixed group’; *n* = 321) comprised athletes whose sports utilized mixed anaerobic and aerobic energy production (6 badminton players, 28 basketball players, 15 water polo players, 29 volleyball players, 26 handball players, 7 curling players, 2 equestrian athletes, 4 freestyle skiers, 13 sailors, 7 divers, 7 lugers, 27 artistic (synchronized) swimmers, 3 skeleton athletes, 16 snowboarders, 10 modern pentathletes, 8 archers, 14 shooters, 23 fencers, 21 figure skaters, 19 field hockey players, 36 ice hockey players). The third group (‘speed-strength group’; *n* = 100) included strength athletes and explosive power athletes with predominantly anaerobic energy production (55 wrestlers, 7 alpine skiers, 2 throwers, 4 jumpers (track and field), 3 heptathletes, 22 artistic gymnasts, 7 weightlifters). The fourth group (‘sprint group’; *n* = 45) included sprinters (2 100-400 m runners, 10 sprint cyclists, 2 200 m kayakers, 6 500-1000 m speed skaters, 19 50-100 m swimmers, 6 500-1000 m short-trackers).

Athletic success was measured by determination of the level of achievement of each athlete. There were 95 athletes classified as ‘highly elite’ (‘merited masters of sport’ according to the unified sports classification system of Russia (USCSR; awards only for results on the official competitions); equates to international champion who has made valuable contributions to the sport (prize winners of Olympic Games and World / Europe championships)). There were 190 athletes classified as ‘elite’ (‘masters of sport of international class’ according to the USCSR; equates to international champions). The other athletes (*n* = 314) were classified as ‘sub-elite’ athletes (‘masters of sport’ (equates to national champions) and ‘candidates for master of sport’ (equates to nationally ranked athletes) according to the USCSR). Controls (*n* = 298) were age-matched (16-35 years old) Russian women without any sports experience.

The procedures followed in the study were conducted ethically according to the principles of the World Medical Association, the Declaration of Helsinki and ethical standards in sport and exercise science research. The Ethics Committee of the All-Russian Research Institute of Physical Culture and Sport approved the study and written informed consent was obtained from each participant.

### Procedures

Resting T (i.e. T which was measured on a day when the athlete did not train) was examined in serum of athletes. For these purposes, 10 ml of venous blood was collected the morning after an overnight fast and sleep into EDTA vacutainer tubes and placed at 4°C until processing. For each sporting discipline blood was collected on the same day at least 15 h after the last training. T was analyzed on a Benchmark Plus Microplate Spectrophotometer (Bio Rad, France) using enzyme immunoassay and commercial test systems (Alkor-Bio, Russia). The athletes were from the same team and trained under supervision of the same coach.

### Statistical analyses

This study was designed to determine the interrelation between T levels and athletic success (coded as 1, 2 and 3 for sub-elite, elite and highly elite, respectively) in a cohort of female athletes participating in different types of sport events. Statistical analyses were conducted using GraphPad InStat (GraphPad Software, Inc., USA). Differences in phenotypes between groups were analysed using unpaired *t* tests. All data are presented as mean (standard deviation, SD). Spearman’s correlation was used to assess the relationship (*r*) between the T level and the level of achievement of each athlete. Athletes with different levels of achievement were also compared between each other using ANOVA. The squared correlation coefficient R^2^ was used as a measure of explained variance. Odds ratio was calculated using Chi-square test. *P* values < 0.05 were considered statistically significant.

## RESULTS

The mean (SD) T levels of athletes and controls were 1.65 (0.87) and 1.76 (0.6) nmol/L (*P* = 0.057 for difference between groups), respectively with range 0.08-5.8 nmol/L in athletes and 0.38-2.83 nmol/L in controls. There were 11.6% female athletes and 7.4% female controls with a T level > 2.7 nmol/L, the upper limit of the normal reference range. T levels slightly increased with the age of all athletes (*P* = 0.139), but significantly decreased with the age in controls (*P* = 0.0042). No significant differences in T levels were found between the different groups of athletes. T levels were positively correlated with athletic success in sprinters (*r*=0.62, *P*<0.0001; *P*_ANOVA_ < 0.0001) only, indicating that 38% of the variation in athletic success in sprinters can be explained by the T levels. None of sub-elite sprinters had T > 1.9 nmol/L, while 50% of elite and highly elite sprinters had T > 1.9 nmol/L (95% CI: 2.562-862.34; OR=47.0; *P* < 0.0001). No correlations were found in other groups of athletes (Table 1).

**Table 1.** Interrelationship between resting serum testosterone levels and athletic success in female athletes stratified into four groups

| Group | n | Testosterone, nmol/L |  |  |  | *r* | *P* value (Spearman) | *P* value (ANOVA) |
| --- | --- | --- | --- | --- | --- | --- | --- | --- |
|  |  | All athletes | Sub-elite | Elite | Highly elite |  |  |  |
| Endurance athletes | 133 | 1.67 (0.85) | 1.68 (0.87) | 1.72 (0.9) | 1.44 (0.58) | 0.064 | 0.467 | 0.556 |
| Athletes with mixed activity | 321 | 1.72 (0.91) | 1.74 (0.99) | 1.81 (0.86) | 1.55 (0.75) | 0.021 | 0.709 | 0.214 |
| Speed/strength athletes | 100 | 1.46 (0.79) | 1.44 (0.78) | 1.47 (0.82) | 1.43 (0.81) | 0.04 | 0.725 | 0.976 |
| Sprinters | 45 | 1.52 (0.73) | 1.09 (0.4) | 1.97 (0.63) | 2.0 (0.93) | 0.62 | <0.0001* | <0.0001* |
| All athletes | 599 | 1.65 (0.87) | 1.64 (0.92) | 1.72 (0.86) | 1.56 (0.75) | -0.006 | 0.881 | 0.337 |

Values are M (SD). \* $P < 0.05$ , statistically significant correlation (Spearman), $P < 0.0001$ (ANOVA).

When endurance athletes were divided into long- and middle-distance athletes, we also could not observe the benefits of having high T levels for success in two types of endurance events, including 800 m running. Instead, highly elite long-distance (1.62 (0.55) vs 1.78 (0.87) nmol/L, *P* = 0.601) and middle-distance (1.09 (0.47) vs 1.57 (0.88) nmol/L, *P* = 0.235) athletes tended to have lower T levels than less successful athletes. The same tendency was observed in the mixed cohort (1.55 (0.75) vs 1.77 (0.95) nmol/L, *P* = 0.096), while there were no differences between highly elite speed/strength athletes and the rest (1.43 (0.81) vs 1.46 (0.80) nmol/L, *P* = 0.931).

## DISCUSSION

Our main finding was that T levels were positively correlated with athletic success in Russian female sprinters. The second main outcome of the study was that T levels did not confer competitive advantage in other types of sport events such as endurance disciplines, events with mixed sport activities and speed-strength sports. Importantly, 11.6% of female athletes from our study had T levels > 2.7 (up to 5.8) nmol/L (above the normal reference range) which is similar (13.7%) with previously published data on 239 female athletes from the UK^21^. The novelty of the study was that the association of the T levels with athletic success was found not only in sprinters specialized in running, but also in swimming, cycling, kayaking, speed skating, and short-track.

Although the role of androgens in female physiology has not been well established, several studies have indicated that T may induce skeletal muscle hypertrophy and increase muscle strength through genomic (long-term, transcriptional) mechanisms^22,23^. Additionally, T was shown to act quickly through non-genomic mechanisms (short-term, non-transcriptional)^24^. Based on the current understanding of non-genomic T action, Dent, Fletcher, McGuigan^25^ suggested that T may be able to produce an increase in intracellular calcium levels and calcium mobility within the human skeletal muscle cell. This may increase the sensitivity of the contractile elements to calcium, which could increase the speed of myosin head binding and/or the force at which the myosin head pulls, such that more force is produced per contraction. Combined, these effects would likely result in greater whole muscle power production and thus, sprinting ability. Consistent with this hypothesis and our data, Cardinale, Stone^22^ identified a significant positive relationship between T levels and vertical jump performance both in male (*r*=0.62) and female (*r*=0.48) elite athletes involved in sprinting, volleyball, soccer and handball. Furthermore, Bosco et al.^26,27^ reported a positive correlation between T levels and vertical jump height and power output in professional male soccer players. The same authors found that T levels were higher in explosive male athletes, such as sprinters, and lowest in endurance athletes such as cross-country skiers^28^. Recently, Bermon et al.^29^ confirmed that T levels of female sprinters were higher in comparison with long distance athletes. It is possible that T might improve athletic performance in sprint events by decreasing reaction time, as T has been shown to regulate neuromuscular transmission^30^. It was, therefore, suggested that T plays an important role in explosive performance by influencing skeletal muscle excitation-contraction coupling, phenotypization of fast-twitch muscle fibers, protein synthesis and aggressive behaviour^22^. Interestingly, in the mixed cohort of 18 female athletes (track and field, netball, cycling, swimming and bob skeleton), Cook, Crewther, Smith^31^ revealed that salivary free T concentrations were significantly higher in elite athletes in comparison with non-elites, suggesting that T could influence the expression of behaviour (i.e. dominance) and thereby help to regulate sporting potential.

We could not take into consideration several possible confounding factors which were not determined, such as the use of oral contraceptives (OC), menstrual status (e.g., oligo- and amenorrhea as well as OC intake leads to lower androgen levels) and some disorders. It’s also well established that a mass spectrometry-based measurement instead of immuno-assay method of T detection should be preferred. These are limitations of this study. In addition, there are already other factors that have been reported to show associations with sprinting ability, such as body mass, body mass index and height^32,33^, muscle fiber composition^34,35^ and several genetic variants^36–41^.

## Conclusions

Taken together, our data indicate that T is responsible for athletic success in female sprinters but not in athletes from other types of sport events such as endurance disciplines, events with mixed sport activities and speed-strength sports. Given that 38% of the variation in athletic success in sprinters can be explained by the T levels, the preliminary data suggest that the measurement of serum T (which is an inherited trait) may be used in the prediction of sprinting ability in women. Additional studies in female athletes from different groups of sport events with T levels above 10 nmol/L (with normal androgen sensitivity and after passing the clinical investigations) are needed to establish whether the hyperandrogenic state gives them competitive advantage. We also propose that current regulations on female hyperandrogenism should be reconsidered and based on the type of sport event.

